# Robust estimation of brain stimulation evoked responses using magnetoencephalography

**DOI:** 10.1101/2023.06.25.546459

**Authors:** Ashwini Oswal, Bahman Abdi-Sargezeh, Tolga Esat Özkurt, Samu Taulu, Nagaraja Sarangmat, Alexander L Green, Vladimir Litvak

## Abstract

Magnetoencephalography (MEG) recordings are often contaminated by interference that can exceed the amplitude of physiological brain activity by several orders of magnitude. Furthermore, activity of interference sources may *‘leak’* into the activity of brain signals of interest, resulting in source estimation inaccuracies. This problem is particularly apparent when using MEG to interrogate the effects of brain stimulation on large scale cortical networks.

This technical report offers two contributions. Firstly, using phantom MEG recordings we describe an approach for validating the estimation accuracy of brain stimulation evoked responses. Secondly, we propose a novel denoising method for suppressing the leakage of stimulation related signal into recorded brain activity. This approach leverages spatial and temporal domain projectors for signal arising from prespecified anatomical regions of interest. We highlight its advantages compared to the benchmark - spatiotemporal signal space separation (tSSS) - and show that it can more accurately reveal brain stimulation evoked responses.

## Introduction

Magnetoencephalography (MEG) is a powerful technique for interrogating brain network dynamics [Baillet, 2017]. Disturbances of network dynamics are increasingly recognised to contribute to the pathophysiology of neurological conditions such as Parkinson’s disease (PD)[Oswal et al., 2013]. MEG has also been used to investigate how therapeutic interventions including brain stimulation techniques can modulate large scale brain networks. Deep Brain Stimulation (DBS) is one example of such a technique that is used to treat PD, which involves electrically stimulating basal ganglia structures such as the subthalamic nucleus (STN) [Limousin and Foltynie, 2019; Lozano et al., 2019].

A significant challenge of MEG is its vulnerability to contamination from physiological and non-physiological artefacts. MEG recordings during brain stimulation are particularly challenging since the magnitude of the stimulation signal far exceeds that of brain activity of interest [Oswal et al., 2016b; Oswal et al., 2016a]. A related problem is that source estimation techniques are ill-posed, meaning that a few hundred channels cannot sufficiently discriminate brain activity in many thousands of voxels. This results in *‘source leakage’*, whereby reconstructions of dipolar sources will be spread over several voxels and temporally correlated in space [Brookes et al., 2012; Colclough et al., 2015]. Importantly, the leakage of stimulation artefacts precludes accurate estimation of the effects of stimulation on a brain region of interest (ROI).

Spatiotemporal signal space separation (tSSS) [Taulu et al., 2004; Taulu and Kajola, 2005; Taulu and Simola, 2006] is a useful approach for suppressing interference sources that are external to the MEG sensor array. This method has two important shortcomings for brain stimulation studies: 1) it ignores the possibility of artefactual sources originating from inside the sensor array, and 2) it does not mitigate against the leakage of stimulation-related activity into the reconstructed activity from brain ROIs [Colclough et al., 2015].

Although tSSS has been used for estimating brain stimulation evoked responses, to our knowledge there have been no reports regarding the accuracy of this approach [Bahners et al., 2023; Hartmann et al., 2018]. Previous work has explored stationary source estimation accuracy in the presence of brain stimulation artefacts [Abbasi et al., 2016; Kandemir et al., 2020; Oswal et al., 2014; Oswal et al., 2016b]. This problem is different for the estimation of evoked responses however, which are short lived and temporally tightly correlated to high amplitude stimulation pulses and subsequent artefacts.

Here we use phantom recordings obtained during DBS to validate the estimation accuracy of evoked responses. Secondly, we develop an extension of the tSSS methodology for anatomical ROIs, to eschew the two shortcomings described above.

## Methods

### Phantom recording for validating estimation accuracy of stimulation evoked responses

We previously described a phantom experiment to characterise DBS artefacts during MEG recordings [Oswal et al., 2016b]. In this, a 27 Hz dipolar source mimics brain activity whilst monopolar DBS is delivered at frequencies of 0 (no stimulation condition), 5, 20 or 130 Hz (pulse width = 60μs; amplitude = 3V). We simulated evoked responses using this data by epoching continuous data segments into trials that were locked to a specific phase of the 27 Hz source.

MEG data - sampled at 2.4kHz using a CTF 275 channel system - were first denoised using either tSSS or our novel method, region of interest based tSSS (ROI-tSSS; see below). A high pass filter (5 Hz) was subsequently applied before epoching. Trial data were averaged, and topographies of the simulated evoked response were compared for the different combinations of stimulation conditions and denoising approaches. A control condition (‘Standard’), where no denoising was applied is also included. Simulated dipole time courses were reconstructed for visualisation using the *ft_dipolefitting* function in FieldTrip [Oostenveld et al., 2011].

### Extending spatiotemporal signal separation (tSSS) for regions of interest (ROI-tSSS)

tSSS uses a spatial filter to decompose an *N* channel MEG signal, *b* into two components (*b*_*in*_ and *b*_*out*_) corresponding to source locations inside and outside the MEG sensor array [Taulu and Simola, 2006].

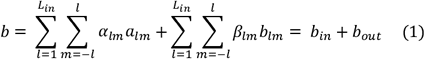

In (1), *a*_*lm*_ and *a*_*lm*_ are SSS basis functions, which depend on sensor geometry and are derived from the gradients of spherical harmonic functions in spherical coordinates. *L*_*in*_ and *L*_*out*_ govern dimensions of the SSS basis set and are prespecified as 8 and 6 respectively [Taulu and Simola, 2006]. Equation (1) expressed in matrix form is:

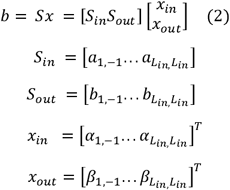

Equation (2) reveals that SSS coefficients, *x*_*in*_ and *x*_*out*_ can be computed from the pseudoinverse of the SSS basis (*S*^*^) set and the data as follows:

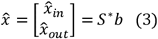

Estimates of *x*_*in*_ and *x*_*out*_ can then be used to estimate *b*_*in*_ and *b*_*out*_:

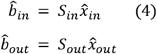

Building on previous results [Ozkurt et al., 2006; Özkurt et al., 2009], we show that manipulating the SSS coefficient *x*_*in*_ can filter the SSS signal into separate components originating from within and from outside a spherical brain ROI (e.g., motor cortex; see **Supplementary Material** for further details). The modified SSS coefficient for the ROI, 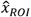 is given by:

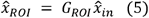

Where:

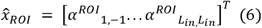

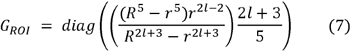

In equation (7) *G*_*ROI*_ is a diagonal matrix, where *l* for each element *G*_*ROI(n,n)*_ is given by 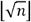: *r* represents the radius of the ROI; and *R* is the radius of the sensor array from the SSS expansion origin, added to the distance between the SSS expansion origin and the centre of the ROI [Özkurt et al., 2009].

Similarly, the modified SSS coefficient for regions outside the ROI, 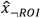 is:

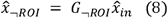

Where *G*_*⌝ROI*_ is a diagonal matrix given by:

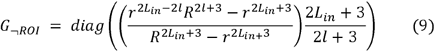

Consequently, estimates of the signal originating from the volume of the ROI 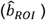 and from the brain volume external to it 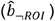 are given by:

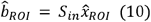

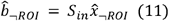

Following the spatial filtering step in tSSS, a temporal filter is applied to remove correlated signal components (representing artefacts) in both 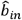 and 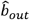 [Taulu and Simola, 2006]. In ROI-tSSS, we apply temporal filtering to remove from 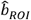 components that are correlated between 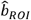 and 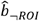 (see **Supplementary Materials**). This procedure removes zero-lag correlated signal components from brain ROI activity and therefore offers source leakage correction.

The above procedure assumes spherical ROIs. In order to achieve improved control of brain volumes encompassed by an ROI, we extended our algorithm to include cuboidal ROIs (see **Supplementary Methods**).

### Application of ROI-tSSS for detecting DBS evoked responses

We applied the ROI-tSSS algorithm to extract cortical evoked responses to 5 Hz monopolar STN DBS in a patient with PD (see **Supplementary Methods**). LCMV beamformer [Van Veen et al., 1997] source localisation was performed for each 5 mm spaced grid point on the 3D brain volume using the SPM DAiSS toolbox (https://github.com/SPM/DAiSS). Data used for beamforming each grid point were processed using the ROI-tSSS algorithm (spherical ROI with 3cm radius), before being high pass filtered (5 Hz) and epoched to the onset of DBS pulses.

## Results

### Phantom recordings: improved estimation of evoked response topographies

Figure 1. shows the topography of the simulated evoked response for the different monopolar DBS frequencies and pre-processing approaches. When no denoising is applied (‘Standard’) the topographies are inaccurately reconstructed across all DBS settings. The ROI-tSSS approach - using either a spherical or cubic ROI centred on the simulated dipole - provides accurate reconstruction of topographies across all stimulation frequencies.

**Figure 1.**
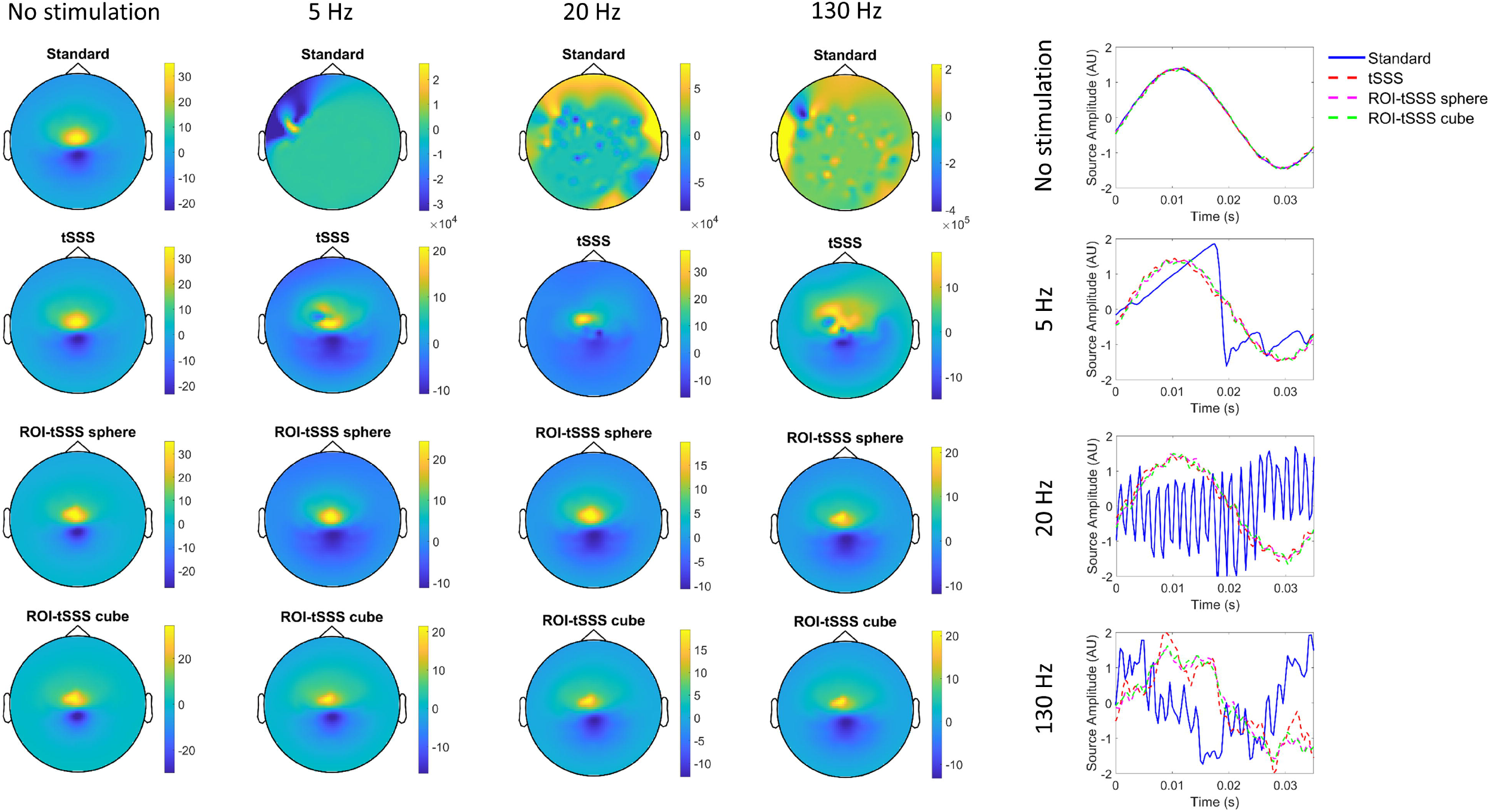
Application of spatiotemporal signal separation approaches to reconstructing evoked responses in a phantom recording. ***Left***: Topographies of the simulated dipole at the four different DBS settings (No stimulation, 5 Hz monopolar DBS, 20 Hz monopolar DBS and 130 Hz monopolar DBS), after pre-processing the data with one of four different approaches (Standard pre-processing, tSSS, ROI-tSSS sphere and ROI-tSSS cube) are shown. The colour bars represent field strength measured in femtoteslas (fT). Plots to the ***right*** show the reconstructed time courses of the simulated sinusoid, for each DBS setting and each pre-processing approach. The ROI-tSSS sphere and ROI-tSSS cube approaches reproduce the dipole topography well across all stimulation conditions.

For each pre-processing approach we computed the mean squared error (MSE) between the sensor timeseries of each stimulation condition and the corresponding no stimulation condition (**Supplementary Figure 1**). ROI-tSSS outperforms tSSS for all stimulation conditions with the difference being particularly pronounced at 130Hz – a commonly employed clinical DBS frequency. **Figure 1** also shows estimated time courses for the simulated dipole for the different DBS conditions and pre-processing approaches.

### Patient recording: Effects of STN DBS on cortical networks

The left panel of ***Figure 2*** shows the mean evoked response, averaged over trials and channels, after constructing a 3cm ROI around the right motor cortex. There are early peaks after 2.5 and 4.2ms. The sensor topographies of these two peaks are also shown, revealing activation of parietal sensors. In the right panel of ***Figure 2***, cortical evoked response amplitudes at the timings of the aforementioned peaks are displayed. In addition to small activations within the precentral gyrus, there are larger amplitude peaks within the middle and inferior frontal gyri, the temporal lobe, and parieto-occipital regions.

**Figure 2.**
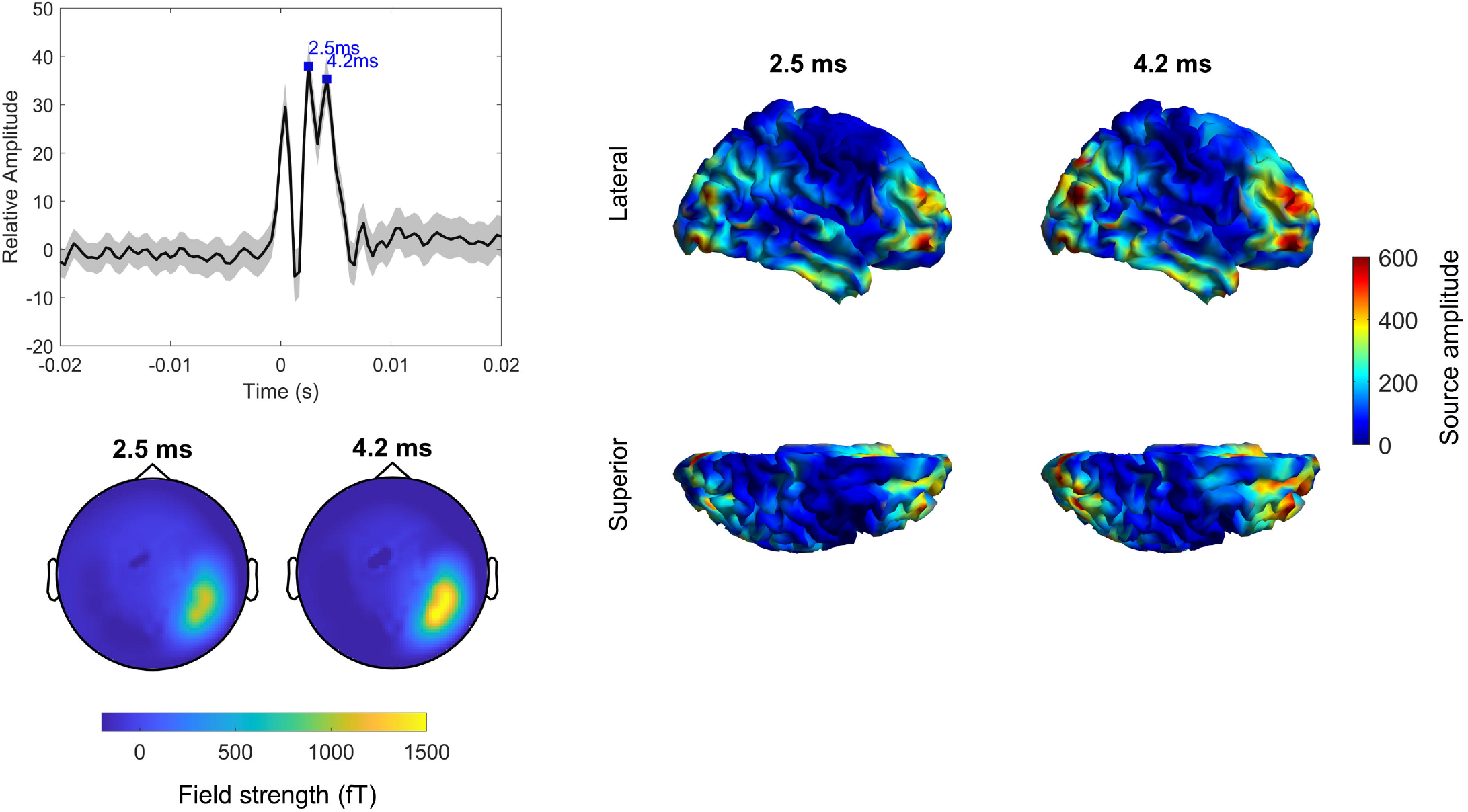
The ROI-tSSS sphere approach is applied to detect the evoked response to monopolar 5Hz DBS of the right subthalamic nucleus in a patient with Parkinson’s disease. ***Left***: upper panel shows the sensor level evoked response (averaged across trials and channels after baseline correcting relative to a 20ms window [-0.05s to -0.03s] prior to the onset of the stimulation pulse at time 0) after constructing a 3cm region of interest centred on the right motor cortex at MNI co-ordinates 37 -18 53; the lower panel shows the corresponding topography of the evoked response peaks at 2.5ms and 4.2ms. ***Right***: right hemispheric source level evoked response amplitudes at the times of the two peaks are extracted using LCMV beamforming and projected onto a cortical mesh. For each grid point, the ROI-tSSS sphere algorithm (with a 3cm radius) was applied prior to beamforming. There are focal frontal, temporal and parieto-occipital regions demonstrating high amplitude evoked responses.

## Discussion

The aim of this report is to validate and advance the use of MEG for characterising the effects of brain stimulation techniques on large scale cortical networks. We have described a phantom recording paradigm for characterising the estimation accuracy of stimulation evoked responses. Secondly, we detail an extension of the tSSS algorithm for suppressing brain stimulation artefacts during MEG recordings. Our approach (ROI-tSSS) involves constructing separate spatial projectors for a brain ROI and for areas outside the ROI. A temporal projector is then used to remove signal components - representing leakage artefacts - that are common to both the ROI and areas outside the ROI.

ROI-tSSS works effectively for reconstructing a simulated evoked response under different DBS conditions in the phantom recording. In recordings from a single PD patient, ROI-tSSS recovered early latency evoked responses to STN DBS from frontal and temporal brain areas. These results closely match the findings of invasive studies using electrocorticography (ECOG) [Chen et al., 2020; Jorge et al., 2022]. We also observed evoked responses within parieto-occipital brain areas - consistent with the existence of functional connectivity networks between these cortical areas and the STN [Litvak et al., 2011].

## Acknowledgements

AO is supported by an MRC Clinician Scientist Fellowship (MR/W024810/1). The Wellcome Centre for Human Neuroimaging is supported by core funding from Wellcome [203147/Z/16/Z].

## Data and code availability

Phantom recording data used in this paper are available at the following repository: https://figshare.com/articles/dataset/phantom090715_BrainampDBS_20150709_01_ds_zip/4042911.

Code for implementing the ROI-tSSS algorithm are shared on GitHub repository https://github.com/AshOswal/ROI-tSSS

---

**Supplementary Figure 1.**
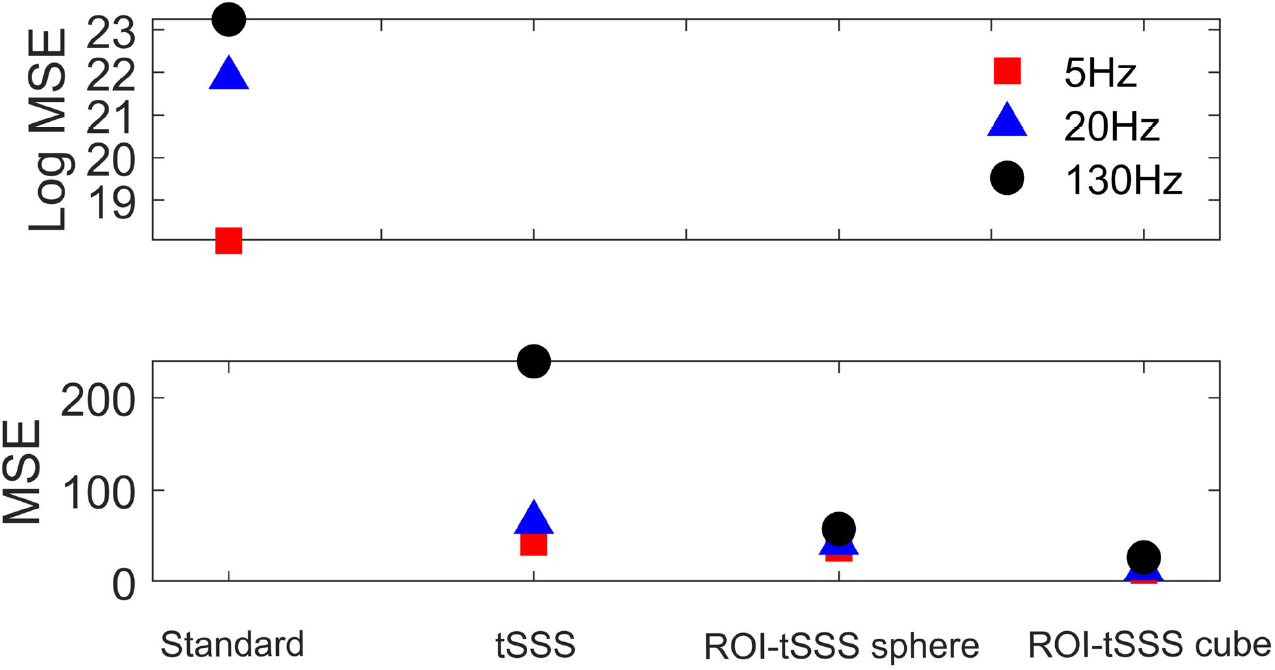

**Supplementary Figure 2.**
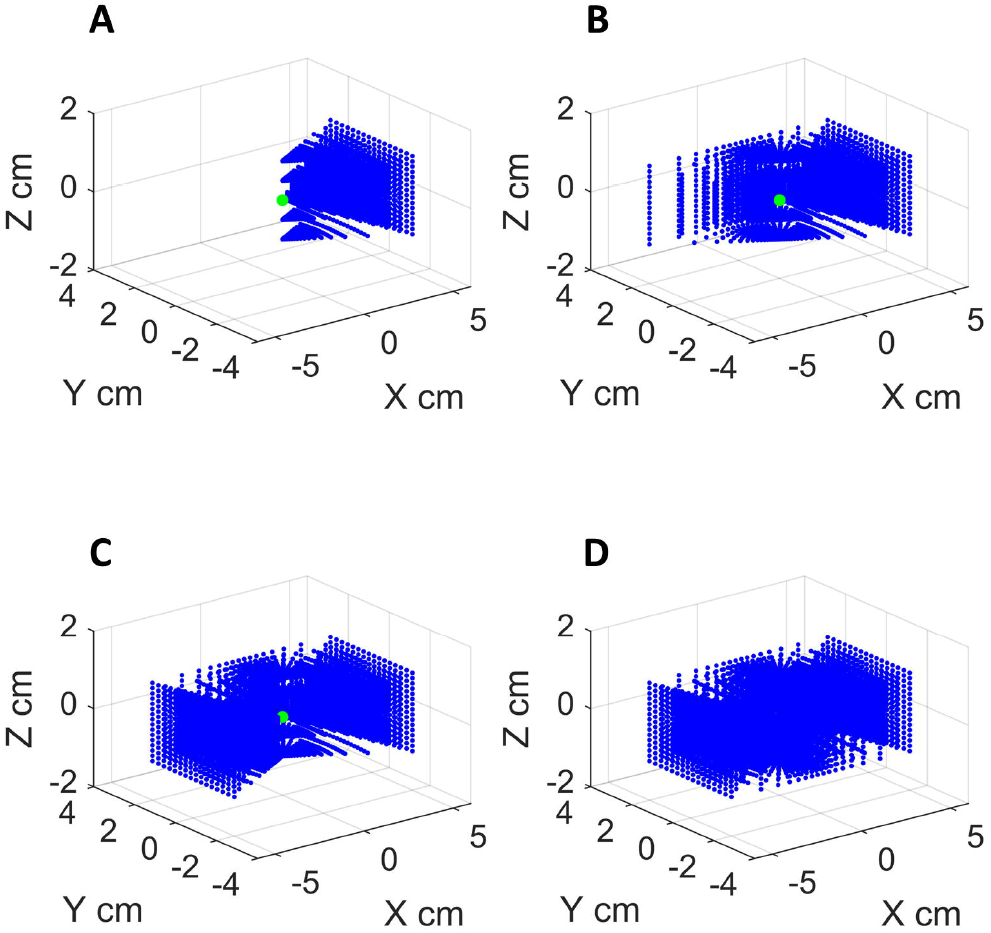

**Supplementary Figure 3.**
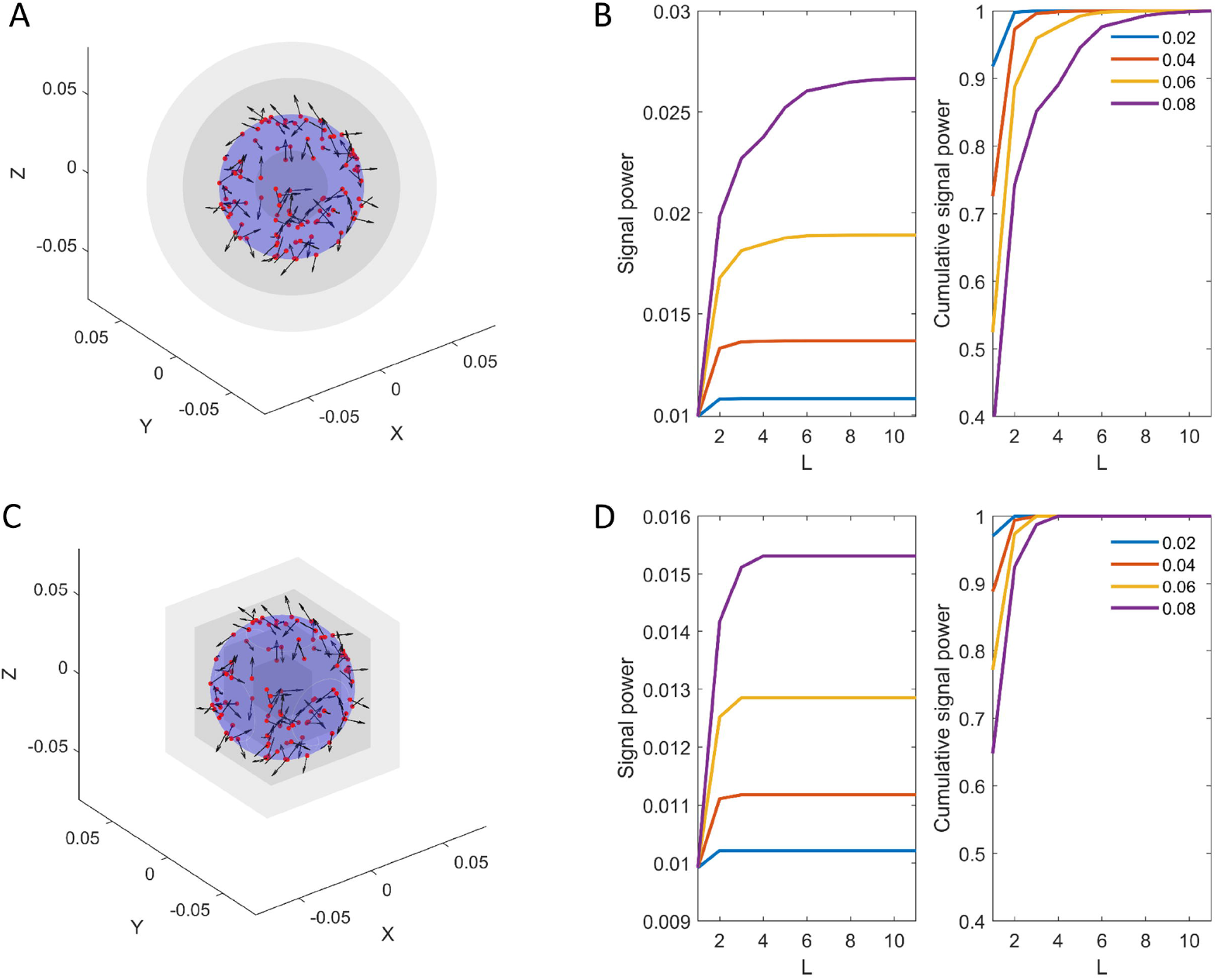

