## Supplementary Methods for "Robust estimation of brain stimulation evoked responses using magnetoencephalography"

**Supplementary Materials**

**Supplementary Methods**

***Derivation of formulae for*** $\boldsymbol{G}_{\boldsymbol{ROI}}$ ***and*** $\boldsymbol{G}_{\boldsymbol{\neg ROI}}$

We first outline how the modified SSS coefficients for the spherical ROI ($G_{ROI}$) and the brain volume outside this ($G_{\neg ROI}$) are constructed.

Previous work has demonstrated that SSS coefficients $(\alpha_{1,-1}..,\alpha_{l,m},... \alpha_{L_{in,}L_{in}})$ with low values of $l$ correspond to sources close to the center of an ROI, whilst those with high values of $l$ correspond to sources that are further away [Ozkurt et al., 2006]. Arbitrarily selecting a cut off value for $l$ can be inaccurate. One way to address this issue is to apply a weighting to the SSS coefficients that is derived by first representing them in a leadfield like manner [Ozkurt et al., 2006; Taulu and Simola, 2006].

$$\alpha_{l,m} = \int\lambda_{l,m}(r).J_{in}(r) dv (1)$$

In (1), $r$ represents the radius of the ROI, $J_{in}(r)$ denotes the current distribution and $\lambda_{l,m}(r)$ is analogous to a conventional leadfield for the SSS coefficient [Taulu and Simola, 2006]. The integral in (1) corresponds to a volume element in spherical coordinates. It can be shown that $\lambda_{l,m}(r)$ is given by [Taulu and Simola, 2006]:

$$\lambda_{l,m}(r)=\frac{i}{2l+1}\sqrt{\frac{l}{l+1}}r^{l}X_{l,m}^{*}(\theta,\varphi) (2)$$

Where $X_{l,m}^{*}$ is a vector spherical harmonic function that is independent of $r$. A Matrix describing second order relationships between the leadfields for the ROI is given by equation (3), where the asterisk represents the complex conjugate:

$$R = \iiint_{r=0}^{r} \left( \frac{i}{2l+1}\sqrt{\frac{l}{l+1}}r^{l}X_{l,m}^{*}(\theta,\varphi) \right)\left( \frac{i}{2l+1}\sqrt{\frac{l}{l+1}}r^{l}X_{l,m}(\theta,\varphi) \right) r^{2}\sin\theta dr d\theta d\varphi(3)$$

This equation can be simplified by first considering the orthonormality property of vector spherical harmonics, where $\delta_{ab}$represents the dirac delta function:

$$\iint X_{l,m}^{*}(\theta,\varphi) .X_{l,m}(\theta,\varphi)\sin\theta d\theta d\varphi= \delta_{lL}\delta_{mM} (4)$$

Integrating over $r$, means that entries $R$ simplify to:

$$R_{ab} = \delta_{ab}\frac{1}{{(2l+1)}^{2}}\frac{l}{l+1}\frac{r^{2l+3}}{2l+3} (5)$$

A similar matrix of the second order relationships between leadfields of regions outside the ROI

Is given as:

$$\neg R = \iiint_{r=r}^{R} \left( \frac{i}{2l+1}\sqrt{\frac{l}{l+1}}r^{l}X_{l,m}^{*}(\theta,\varphi) \right)\left( \frac{i}{2l+1}\sqrt{\frac{l}{l+1}}r^{l}X_{l,m}^{*}(\theta,\varphi) \right) r^{2}\sin\theta dr d\theta d\varphi(6)$$

$R$ is the radius of the sensor array from the SSS expansion origin, added to the distance between the SSS expansion origin and the centre of the ROI. Again, using orthonormality properties, (5) simplifies to:

$${\neg R}_{ab} = \delta_{ab}\frac{1}{{(2l+1)}^{2}}\frac{l}{l+1}\frac{R^{2l+3} - r^{2l+3}}{2l+3} (7)$$

Ozkurt et al. [Özkurt et al., 2009] show that the power of the sources within the ROI relative to source power outside the ROI can be maximized by constructing a weighting for the SSS coefficients, $G$:

$$G = \frac{v_{roi}}{v_{\neg roi}}{\neg R}^{-1/2}R{\neg R}^{-1/2} = \left( \frac{{(R}^{3}-r^{3})r^{2l}}{R^{2l+3}-r^{2l+3}} \right) (8)$$

Where $v_{roi}$ represents the volume of the ROI and $v_{\neg roi}$ represents the volume outside the ROI. Similarly, the weighting of SSS coefficients for areas outside the ROI is given by the reciprocal of $G$:

$$\neg G = \frac{v_{\neg roi}}{v_{roi}}R^{-1/2}\neg RR^{-1/2} = \left( \frac{R^{2l+3}-r^{2l+3}}{{(R}^{3}-r^{3})r^{2l}} \right) (9)$$

Note that both $G$ and $\neg G$ are diagonal matrices. We prefer to normalise the entries, by dividing by the largest value (or multiplying by the reciprocal of this), so as to avoid very small or very large scaling factors (see Figure 3 in [Ozkurt et al., 2006]). This gives:

$$G = \left( \frac{{(R}^{3}-r^{3})r^{2l}}{R^{2l+3}-r^{2l+3}} \right) \left( \frac{R^{5}-r^{5}}{{(R}^{3}-r^{3})r^{2}} \right) \to\left( \frac{{(R}^{5}-r^{5})r^{2l-2}}{R^{2l+3}-r^{2l+3}} \right) (10)$$

$$\neg G = = \left( \frac{R^{2l+3}-r^{2l+3}}{{(R}^{3}-r^{3})r^{2l}} \right) \left( \frac{{(R}^{3}-r^{3})r^{2L_{in}}}{R^{2L_{in}+3}-r^{2L_{in}+3}} \right) \to\left( \frac{{r^{2L_{in}-2l}R}^{2l+3}-r^{2L_{in}+3}}{R^{2L_{in}+3}-r^{{2L}_{in+}3}} \right) (11)$$

Finally, for both $G$ and $\neg G$ we use L'Hôpital's rule to derive limits for the case when $r =R.$ These limits are then used to normalise the entries of $G$ and $\neg G$ as per [Ozkurt et al., 2006].

$$\lim_{r\to R} \left( \frac{{(R}^{5}-r^{5})r^{2l-2}}{R^{2l+3}-r^{2l+3}} \right) = \frac{5}{2l+3} (12)$$

$$\lim_{r\to R} \left( \frac{{r^{2L_{in}-2l}R}^{2l+3}-r^{2L_{in}+3}}{R^{2L_{in}+3}-r^{{2L}_{in+}3}} \right) = \frac{2l+3}{2L_{in}+3} (13)$$

Using these terms, the final weightings of the SSS coefficients for the ROI and for the volume outside the ROI are given by:

$$G_{ROI} = diag\left( \left( \frac{{(R}^{5}-r^{5})r^{2l-2}}{R^{2l+3}-r^{2l+3}} \right)\frac{2l+3}{5} \right) (14)$$

$$G_{\neg ROI} = diag\left( \left( \frac{{r^{2L_{in}-2l}R}^{2l+3}-r^{2L_{in}+3}}{R^{2L_{in}+3}-r^{{2L}_{in+}3}} \right)\frac{2L_{in}+3}{2l+3} \right) (15)$$

***Computing*** $\boldsymbol{G}_{\boldsymbol{ROI}}$ ***and*** $\boldsymbol{G}_{\boldsymbol{\neg ROI}}$ ***for non-spherical ROIs***

The approach described in the previous section applies to spherical ROIs. In this section we also consider the possibility of deriving projectors for non-spherical ROIs. For instance, for imaging long and thin structures (e.g., the hippocampus) a cuboidal ROI may be more appropriate.

Extension to non-spherical ROIs is challenging since the integrals above in equations (3) and (6) are performed in spherical coordinates and by virtue of this leverage orthonormality properties of vector spherical harmonics. Nevertheless, it is possible to modify the bounds of the integral so that the integration volume covers a cuboid rather than a sphere. This procedure can be performed numerically using a triple integrator (we used the MATLAB function *integral3.m*). We provide a code example of how the integration bounds can be altered to cover a cuboid – see the function *cuboidal_integral_sph.m*. In this function we use these bounds to compute the volume integral of a cuboid in spherical co-ordinates. This involves splitting the cuboid into four triangular prisms and defining integration bounds separately for each of these (**see Supplementary Figure 2**).

For the phantom experiment we compared the ROI-tSSS algorithm for both spherical and cubic ROIs (note that a cube is a special case of a cuboid where all sides are equal). For spherical ROIs we used a radius of 3cm. For the cubic ROI we used a cube with sides that gave a body diagonal of 3cm.

***Construction of power functions***

It has previously been shown [Taulu and Kajola, 2005] that the dependence of signal power recovery on $l$ is given by:

$$\sqrt{\sum_{m} f_{l,m}(r_{\alpha},R,J_{in})} (16)$$

Where:

$$f_{l,m}(r_{\alpha},R,J_{in}) = \sqrt{\frac{l}{2l+1}}\iiint_{0}^{R} \left( \frac{r_{\alpha}}{R} \right)^{l+2}{iX}_{l,m}^{*}(\theta,\varphi). J_{in}(r_{\alpha}) dr d\theta d\varphi(17)$$

In equation (13) $R$ represents the radial distance of the sensor array from the expansion origin, whilst $r_{\alpha}$ is the radius of the source volume. This expansion can be used to determine the optimal value for $L_{in}$ (see Figure 1 in [Taulu and Simola, 2006]). We modify this function for regions of interest by scaling by $G_{ROI}(r)$*,* which is a function of the ROI radius, $r$.

$$f_{l,m}^{ROI}({r, r}_{\alpha},R,J_{in}) = G_{ROI}(r)\sqrt{\frac{l}{2l+1}}\iiint_{0}^{R} \left( \frac{r_{\alpha}}{R} \right)^{l+2}{iX}_{l,m}^{*}(\theta,\varphi). J_{in}(r_{\alpha}) dr d\theta d\varphi(18)$$

This modified power function allows us to consider the relationship between and signal power recovery, $r$ and $l.$ We simulated 100 randomly oriented current dipoles on the surface of a 4cm ($r_{\alpha}$) sphere, whilst $R$ was fixed to 8cm. The function $\sqrt{\sum_{m} {f_{l,m}^{ROI}f}_{l,m}(r,r_{\alpha},R,J_{in})}$ was then computed for different values of $r$ and $l$. The results of this simulation are shown in **Supplementary Figure 3** for spherical and cubic ROIs (for cubic ROIs we selected dimensions such that the cube fitted perfectly within a sphere of a particular radius). When the ROI does not encompass simulated sources, little signal power is recovered as expected. Signal power recovery increases as $r \to R$. To preserve the mapping between sensor space and source space representations when performing source analysis, we applied the spatial component of the ROI-tSSS filter to the leadfields using the *spm_eeg_montage* function.

***Removing zero-lag temporally correlated signal (corresponding to leakage) from*** ${\hat{\boldsymbol{b}}}_{\boldsymbol{ROI}}$ ***and*** ${\hat{\boldsymbol{b}}}_{\boldsymbol{\neg}\boldsymbol{ROI}}$

We construct a projector to remove components that are temporally correlated between the signal estimate from the ROI ($\hat{b}_{ROI}$) and the signal estimate representing regions outside of the ROI ($\hat{b}_{\neg ROI}$). Here we provide a brief overview of this technique, but further mathematical details can be found elsewhere [Taulu and Simola, 2006].

Although this approach is mathematically equivalent to the approach used in tSSS, an important distinction is that it is applied to data representing distinct brain sources or regions. Consequently, it is similar to leakage correction routines based on orthogonalization or regression, which remove signal - representing source leakage - that is correlated at zero-lag [Brookes et al., 2012; Colclough et al., 2015]. Unlike traditional leakage correction techniques that are applied after source analysis, our approach can be applied to sensor data.

The first step involves separately constructing temporal basis sets for both $\hat{b}_{ROI}$and $\hat{b}_{\neg ROI}$, by taking the right singular vectors of their singular value decompositions (SVD). We denote these bases $V_{\hat{b}_{ROI}}$and $V_{\hat{b}_{\neg ROI}}$ respectively. The intersection of the two bases can be determined by computing the SVD of their product.

$$V_{\hat{b}_{ROI}}^{T}V_{\hat{b}_{\neg ROI}} = {USJ}^{T} (19)$$

In this case the singular values (in the diagonal of $S$) contain the cosine of the principal angles between corresponding columns of the two basis sets. In the case of orthogonality, the principal angle is 90 degrees, corresponding to a cosine of 0. Similarly, if the principal angle is 0, corresponding to a cosine of 1, the two basis vectors intersect. Typically, a threshold of between 0.95 and 1 is used to select the number of columns of the matrix $V_{\hat{b}_{ROI}}^{T}U$, which defines the intersection subspace. We call the matrix formed from the first $r$ columns of $V_{\hat{b}_{ROI}}^{T}U$, $R$.

The final ROI-tSSS signal $\hat{b}_{tROI}$ can therefore be obtained by multiplying the ROI signal ($\hat{b}_{ROI}$) onto the subspace orthogonal to the intersection subspace.

$$\hat{b}_{tROI}= \hat{b}_{ROI}(I-RR^{T}) = \hat{b}_{ROI}P (20)$$

***Details of patient recording***

We present data from a 43-year-old PD patient undergoing bilateral STN DBS. A Medtronic 3389 lead was bilaterally implanted, and electrode locations were confirmed with intraoperative MRI (fast spin-echo T2-weighted sequence). Stainless steel electrode extension cables were externalized through the scalp to enable recordings prior to connection to a subcutaneous DBS pacemaker, implanted in a second operative procedure seven days later.

MEG recordings were performed using a CTF 275 channel system, with a sampling frequency of 2.4kHz. A Medtronic external stimulator was used to deliver unilateral monopolar 5Hz DBS (pulse width = 60 µs; amplitude = 3 V) between contact 1 of the DBS electrode (cathode) and an anodal electrode applied to the patient’s chest. at a frequency of 5Hz. Further details regarding the operative procedure and recordings can be found in our previously published work [Oswal et al., 2016]. Recordings were approved by the National Research Ethics Service Committee South Central – Oxford B, and the patients gave written informed consent prior to participation.

**Supplementary Figure Legends**

**Supplementary Figure 1.** The mean squared error (MSE) of the simulated dipole topography, computed between different DBS conditions and the no stimulation condition is plotted for all four pre-processing approaches in the phantom recording. The lowest MSE across all stimulation conditions is achieved by the ROI-tSSS sphere and the ROI-tSSS cube approaches. The ROI-tSSS approaches offer greatest benefit at clinically deployed DBS frequencies of 130 Hz.

**Supplementary Figure 2.** The ROI-tSSS method is extended to cuboidal rather than purely spherical ROIs. Here we consider the integration bounds of a cuboidal ROI of x, y and z dimensions 10cm, 4cm and 2cm. The shape can be split into two pairs of identical prisms, which are serially added moving from A through to D, giving rise to the total volume. Regularly spaced coordinates selected along the integration bounds $(r,\varphi,\theta)$ in the spherical coordinate system are transformed into cartesian coordinates for plotting. The green point indicates the origin (0,0,0).

**Supplementary Figure 3.** The effect of ROI size on signal recovery for spherical and cubic ROIs. ***Upper panel***: (A) highlights the setup of the simulation with 100 randomly oriented sources on the surface of the blue sphere at a distance of 0.04m from the expansion origin. The sensor distance from the origin, *R* is fixed at 0.08m. Grey spheres represent spherical boundaries of the ROIs that were tested (B) the effect of ROI radius on signal power and cumulative signal power. When the ROI does not encompass the simulated sources a small proportion of the signal power is recovered and there is little dependence on L. Signal power recovery and the dependence on L increase as the ROI radius approaches *R.* ***Lower panel*** (C & D) is as per upper panel, except for a cubic rather than a spherical ROI. For cubic ROIs, we selected the dimensions of the side such that the body diagonal was equal to the diameter of the corresponding sphere in (A).
